# AXL limits the mobilization of cholesterol to regulate dendritic cell maturation and the immunogenic response to cancer

**DOI:** 10.1101/2023.12.25.573303

**Authors:** Meriem Belabed, Matthew D. Park, Cédric M. Blouin, Sreekumar Balan, Chang Y. Moon, Jesse Boumelha, Ante Peros, Raphaël Mattiuz, Amanda M. Reid, Camillia S. Azimi, Nelson M. LaMarche, Leanna Troncoso, Angelo Amabile, Jessica Le Berichel, Steven T. Chen, C. Matthias Wilk, Brian D. Brown, Kristen Radford, Sourav Ghosh, Carla V. Rothlin, Laurent Yvan-Charvet, Thomas U. Marron, Daniel J. Puleston, Nina Bhardwaj, Christophe Lamaze, Miriam Merad

## Abstract

We previously found that uptake of cellular debris prompts conventional dendritic cells (cDCs) to undergo maturation. This transformation results in DCs entering the molecular state termed ‘mregDC’. In this state, mregDCs dampen their ability to acquire new antigens, upregulate chemokine receptors to migrate to lymphoid organs, and upregulate MHC-I and -II, co-stimulatory, and -inhibitory molecules to promote the differentiation of antigen-specific T cells. Here, we show that cholesterol mobilization – through both *de novo* synthesis and the acquisition of the metabolite during debris uptake – drives cDCs to mature into mregDCs. This cholesterol is used to assemble lipid nanodomains on the plasma membrane of mregDCs to support cell surface expression of maturation markers. This process is dependent on both *de novo* synthesis and Niemann-Pick disease type C1 (NPC1), which shuttles cholesterol from the endolysosomal pathway. Specifically, NPC1 mediated the accumulation of IFN-ɣ receptor (IFNɣR) in cell surface lipid nanodomains, enabling optimal IFNɣR signaling required for IL-12 production and efficient T cell activation. Importantly, we also show that the receptor tyrosine kinase AXL constitutively dampens the cholesterol-dependent construction of lipid nanodomains on mregDCs; its deletion from cDCs enhance mregDC immunogenicity and yielded potent anti-tumor immunity in an experimental model of lung cancer. Altogether, our findings present novel insights into the mobilization of cholesterol for proper immune receptor signaling as a basis for cDC maturation and the novel role of AXL as a central regulator of this process that can be therapeutically targeted to leverage the immunostimulatory features of mregDCs.

## Main Text

Resting conventional dendritic cells (cDCs) in non-lymphoid tissues bear significant phagocytic potential but are considered functionally immature, considering their limited ability to activate T cell programs ^23,35,52^. In contrast, cDCs that have migrated to draining lymph nodes (dLNs), following uptake of antigen, have undergone a notable molecular transformation that enables their antigen-presenting cell function. This occurs at both the steady-state and inflamed setting and involves the co-expression of (i) cytoskeletal machinery that controls cDC migration to T cell zones and adhesion molecules to support their physical interaction with T cells, (ii) elevated expression of MHC class I and II and co-stimulatory ligands that promote antigen presentation and activation of T cells, (iii) inhibitory checkpoints and pro-apoptotic molecules that deter potentially damaging autoinflammation, and (iv) cytokines that instruct T cell differentiation ^17,31,32,41,42^. Earlier technologies have lacked the resolution that is available today, so cells having undergone this change were previously identified simply as CCR7-expressing cDCs that migrated to the dLNs. This cell state was thought to be driven by stochastic migratory cues, motivating the labeling of these cells as ‘migratory cDCs’ ^7,22,35^. Contemporary single-cell technologies, though, have been able to identify CCR7^+^ cDCs in non-lymphoid tissues and among genetically modified cDCs rendered unable to respond to migratory cues ^28^, thus challenging the relevance of the ‘migratory cDC’ nomenclature.

We subsequently found that the capture of cell debris – independently of migratory cues – drives the molecular transformation of cDCs into the ‘migratory’ state, unraveling for the first time the contribution of cellular uptake in dictating cDC maturity ^28^. Yet, capture of cell debris is a process central to the cDC lifecycle, both at steady-state and in disease, influencing their ability to regulate or activate a T cell response. Therefore, understanding the molecular changes that occur upon uptake of cell-associated antigen cargo is imperative, as it reflects a major gap in our study of the regulators of tolerance versus autoimmunity.

Feeding primary cDCs with apoptotic cells *in vitro* drives them to mature, based on the increased expression of activating (CD40, CD80, CD86, MHCI , MHCII, IFN-ɣR) and inhibitory markers like PD-L1 and PD-L2 (**Extended Data Fig. 1a-f**) ^28^. Extensive, orthogonal studies showed that the gene program encoding this transformation is concordant with the signature that defines migratory cDCs ^35^. As such, because this program could also be identified among non-migratory cDCs located in non-lymphoid tissues, we reasoned that a more apt term should reflect the genes belonging to this program. Therefore, we coined the term ‘mature DCs enriched in immuno-regulatory molecules’ or ‘mregDC’, as it represented the co-expression of both immuno-stimulatory and -inhibitory genes within individual cDCs that had matured. Since then, scRNAseq of cDCs from patients identified the mregDC state amongst cDCs isolated from multiple human non-lymphoid tissues in the steady-state and across various disease conditions ^28,37^.

To specify an mregDC signature that is conserved across different non-lymphoid tissues, we performed our own independent analysis of mregDCs by collating publicly available scRNAseq data on human and murine cDCs ^3,5,24,27,40,51,57^ (**Fig. 1a**). These datasets sampled multiple tissues (pancreas, lung, breast, liver, and colon) and varying disease conditions, including healthy specimens. For each species, study-defined cDC clusters were subset from each dataset, integrated in a batch-aware manner, and re-clustered. In addition, highly variable gene sets (hvgs) were identified, and a shared library of homologous genes between the human and murine transcriptomes was then generated and sub-modules of highly correlated hvgs were computed, as previously described ^24,30^. A group of eleven sub-modules constituted a broader gene program, comprised of genes co-enriched with CCR7, the canonical marker for mregDCs (**Supplementary Table 1**). Projection of the mregDC signature onto the integrated re-clustering identified cells in the mregDC state. As these cells were represented in all datasets (**Supplementary Table 2**), we, therefore, concluded that this cell state is not limited to the lungs and confirmed that it is acquired independent from physical migration to dLNs. Therefore, we sought to leverage this integrated analysis to interrogate underlying, conserved mechanisms for cDCs maturing into the mregDC state.

**Fig. 1.**
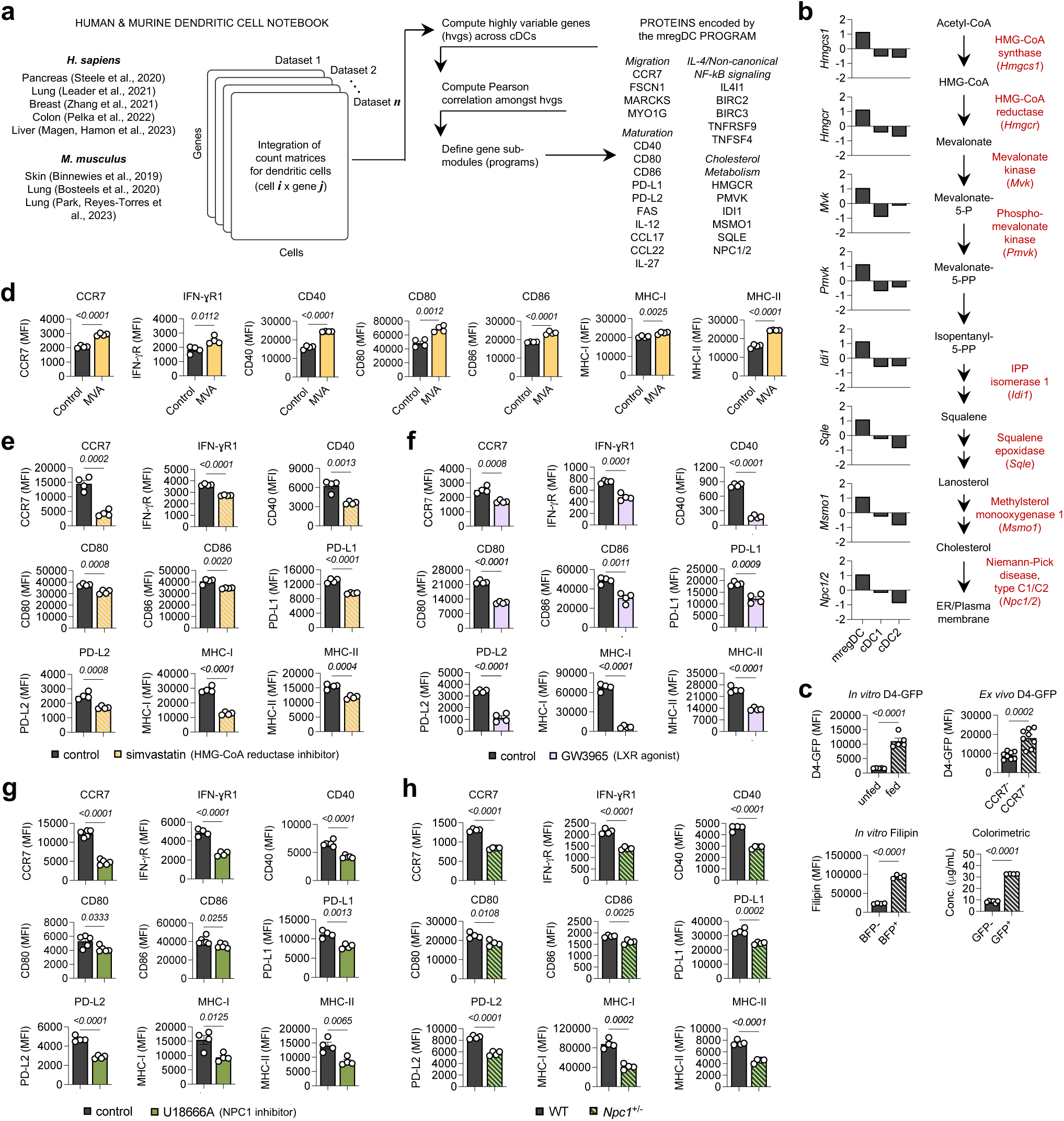
A cellular reservoir of free cholesterol is essential for dendritic cell maturation. (**a**) Computational analysis pipeline for the integrated study of conventional dendritic cells (cDCs) from different tissues across different pathophysiological conditions to identify an mRNA program of cDC maturation. (**b**) Expression (*z*-score) of genes belonging to the cholesterol metabolism signature of the mregDC program. (**c**) Quantification of free cholesterol based on flow cytometric measurement of signal from the D4-GFP probe used to stain DCs *in vitro*, from Filipin dye applied to cDCs (BFP^-^) or to mregDCs (BFP^+^) *in vitro*, colorimetric signal, or from the D4-GFP probe used to stain cDCs and mregDCs *ex vivo* from tumor-bearing lungs. Flow cytometric measurement of cDC maturation markers (i.e., CCR7, IFN-ɣR, CD40, CD80, CD86, PD-L1, PD-L2 or MHC-I, MHC-II) on splenic mregDCs either treated with (**d**) supplemental mevalonate, (**e**) simvastatin, (**f**) the LXR⍺/β agonist GW3965, (**g**) the NPC1 inhibitor U18666A, and (**h**) wild-type (WT) and *Npc1* haploinsufficient *(Npc1*^+/-^) mregDCs. Across all panels, data represent mean ± SEM. *P*-values computed by unpaired *t-*test.

In line with its initial description from single-cell profiling of human and murine cDCs from lung tissues, the reproduction of the mregDC program in this analysis revealed multiple families of genes encoding unique functions (i.e., migration, maturation, IL-4 signaling, and non-canonical NF-κB signaling) (**Fig. 1a**). Notably, we found a family of genes within the mregDC program, encoding enzymes involved in cholesterol metabolism (**Fig. 1b**). As the cholesterol gene network was identified as one of the most significant metabolic pathways (*p*-value = 0.00429), we hypothesized that this metabolic shift may contribute significantly to the acquisition of the mregDC state (**Supplementary Table 3**). Expression of genes belonging to this network (*Hmgcs1, Hmgcr, Mvk, Pmvk, Idi1, Sqle*, *Msmo1*, and *Npc1* and *Npc2*) were enriched in mregDCs, compared to cDCs (**Fig. 1b**); these genes encode enzymes involved in the synthesis of cholesterol via the mevalonate intermediate and transporters like Niemann-Pick disease type C1/2 (NPC1/2) that traffic cholesterol to the plasma membrane ^53^. We confirmed this transcriptional signature by analyzing sorted cDCs and mregDCs that were characterized with bulk RNA sequencing (**Extended Data Fig. 1g,h**) ^28, 64^.

To experimentally validate the role of this cholesterol pathway in mregDCs, we first sought to determine whether cholesterol enrichment is a feature of mregDCs. To test this, we leveraged our reductionist model for efferocytosis, where cDCs are fed with apoptotic blue fluorescent protein (BFP^+^)-tagged cargo derived from ultraviolet (UV)-irradiated cells expressing BFP as a surrogate antigen. In addition to using the polyene macrolide Filipin and an orthogonal colorimetric assay for free cholesterol, we used a green fluorescent protein-tagged probe (D4-GFP) – a fusion of GFP and the D4 fragment of perfringolysin O, a pore-forming toxin from *Clostridium perfringens*, that specifically binds cholesterol ^39,50,55^ – to label and quantify cholesterol in the cell membrane of mregDCs, compared to immature cDCs. Using this tool, we measured GFP signal in BFP^+^ and BFP^-^ DCs from *in vitro* cultures. We also analyzed sorted CCR7^-^ and CCR7^+^ DCs that were isolated from digested tumor-bearing lungs, as this enabled us to look at a greater number of mregDCs *in vivo* than in the steady-state setting. In both comparisons, we found a significant enrichment for cholesterol in mregDCs. By flow cytometry, we also observed significantly higher levels of cholesterol in BFP^+^ cDCs, suggesting that cholesterol does accumulate following uptake – in line with the scRNAseq data (**Fig. 1c**).

Given this result, we perturbed different components of cholesterol metabolism, including synthesis, efflux, trafficking, to determine the role of this internal reservoir of cholesterol in the maturation of cDCs into the mregDC state. Using our *in vitro* culture of cDCs, we first assessed the specific effect of supplementing these cells with exogenous, supraphysiological levels of mevalonic acid (MVA) on the expression of immunogenic markers, namely CCR7; the IFN-ɣ receptor; co-stimulation molecules CD40, CD80, and CD86; and MHC classes I and II. This metabolite drives the *de novo* synthesis of cholesterol, and expression of all markers was significantly elevated on cDCs treated with MVA (**Fig. 1d**). Conversely, blocking cholesterol synthesis with simvastatin (i.e., inhibiting HMG-CoA reductase) or promoting cholesterol efflux by agonizing the liver X receptor (LXR) pathway, both of which would diminish the internal cholesterol reservoir of mregDCs, significantly reduced the cell surface expression of both the immunostimulatory and inhibitory markers associated with maturation. However, for cholesterol-independent markers like the transferrin receptor (CD71), which is known to depend on actin instead for its localization at the plasma membrane ^58^, we found that reducing the intracellular reservoir of cholesterol increased its expression, consistent with published studies on the actin reorganization that accompanies cholesterol depletion ^65^ (**Fig. 1e,f**, **Extended Data Fig. 1i**). Altogether, these data suggest that an enhanced cholesterol reservoir within cDCs contributes to their transformation into mature DCs.

As the expression of the transporters NPC1/2 was highest among genes, we reasoned that properly maturating into mregDCs may be further dependent on the intracellular trafficking of cholesterol *via* these transporters to ensure the optimal allocation of cholesterol. As cholesterol is transferred from endosomal vesicular carriers to the plasma membrane – and given the previously demonstrated link between the mregDC state and efferocytosis – we postulated that the mregDC state is critically dependent on the intracellular transport of cholesterol during efferocytosis, as it is a fundamental step shared by both the *de novo* synthesis of cholesterol and the re-packaging of imported cholesterol. To this end, we asked whether targeting NPC1 with the widely used NPC1 inhibitor (U18666A) ^59^ modulates the expression of maturation markers, compared to those treated with a DMSO control. Strikingly, NPC1 inhibition of mregDCs significantly reduced the cell surface expression of maturation markers, but not the transferrin receptor (**Fig. 1g**, **Extended Data Fig. 1i**). Accordingly, mregDCs generated from mice lacking a single allele for the *Npc1* gene (*Npc1*^+/-^) also exhibited impaired expression of maturation markers (**Fig. 1h**). Altogether, these results demonstrated the importance of a robust cellular reservoir of mobilized cholesterol in defining the mregDC state.

As many of the cDC maturation markers – like the IFN-ɣR and MHC-II – are known to rely on lipid nanodomain structures on the plasma membrane for signaling and function, we hypothesized that the cholesterol mobilized in mregDCs may be used to construct these lipid nanodomains. Using the cholera toxin B subunit (CTxB) to mark lipid nanodomains based on its recognition of the GM1 ganglioside enriched in these structures ^60,61^, we showed that mregDCs exhibit a significantly higher enrichment for lipid nanodomains on the plasma membrane, compared to cDCs, both in *in vitro* culture and *ex vivo* analyses of tumor-bearing lungs (**Fig. 2a**,**b**). CTxB staining was significantly higher on both cDCs supplemented with mevalonic acid and mregDCs but was significantly reduced when the internal cholesterol reservoir was depleted (i.e., treatment with simvastatin or the LXR agonist GW3965) or when cholesterol trafficking was immobilized (i.e., treatment with the NPC1 inhibitor U18666A (**Fig. 2c**,e, **Extended Data Fig. 2a, b**). Importantly, lipid nanodomains were improperly assembled at the cell surface of mregDCs when cells were treated with either methyl-β-cyclodextrin, which depletes all cholesterol from the plasma membrane, or the hydrolase sphingomyelinase, which specifically removes cholesterol sequestered with sphingomyelin (i.e., lipid nanodomains), suggesting that the mobilized cholesterol is essential for the proper assembly of lipid nanodomains (**Fig. 2d,e**). Measurement of free cholesterol with the D4-GFP supported these observations (**Extended Data Fig. 2c-e**). Therefore, when we genetically disrupt cholesterol trafficking using *Npc1*^+/-^ DCs, we could confirm that even a single allelic loss impaired cholesterol mobilization to the plasma membrane and subsequent lipid nanodomain assembly (**Fig. 2f**). Importantly, we found that these lipid nanodomains are indeed essential for the cell surface expression of maturation markers, as treatment with either methyl-β-cyclodextrin or sphingomyelinase abrogated their expression on the plasma membrane (**Fig. 2g,h**), without impacting other cell surface receptors that operate independently of cholesterol, like the transferrin receptor (**Fig. 2i**).

**Fig. 2.**
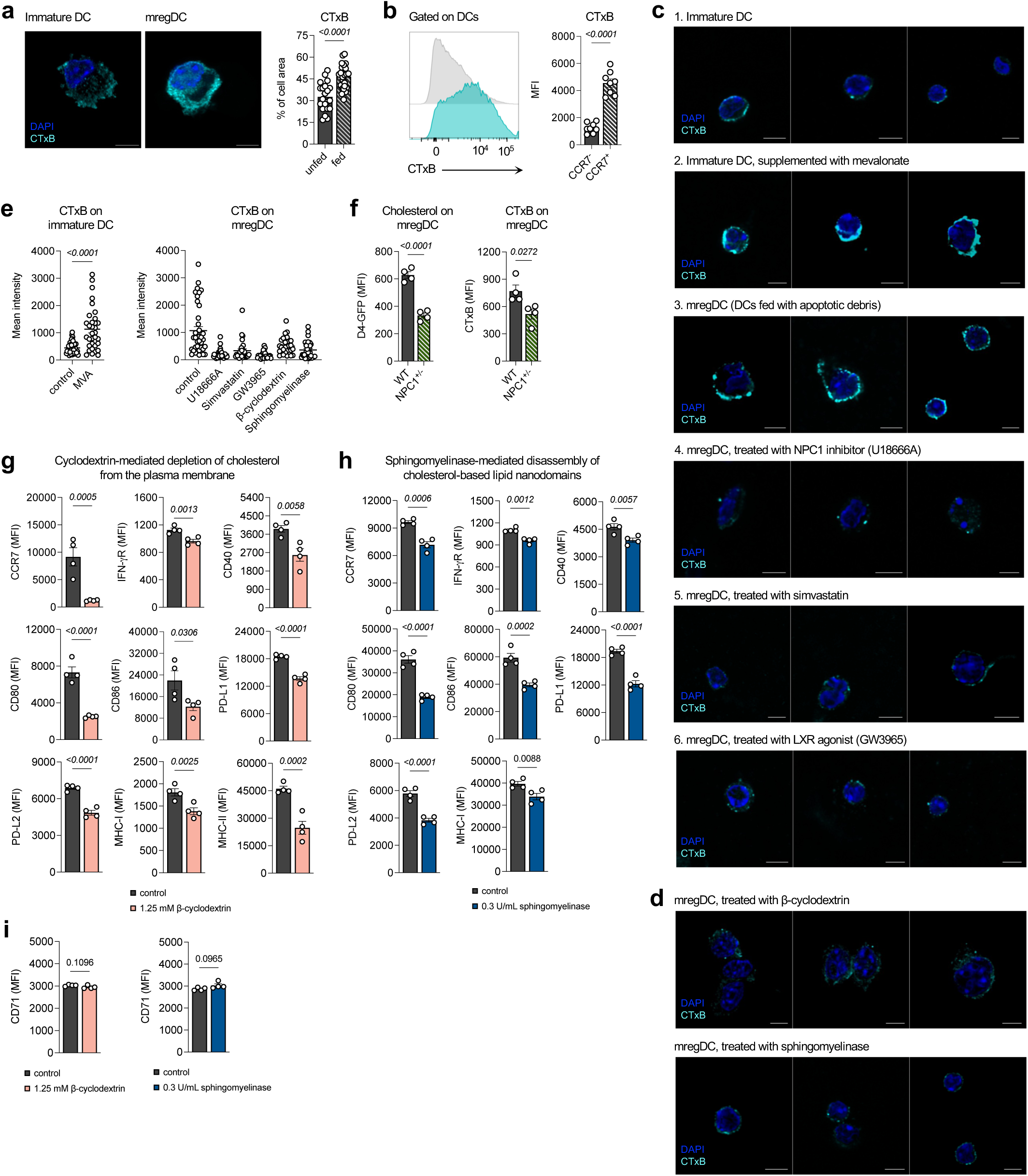
Cholesterol is mobilized to construct cell surface lipid nanodomains that are essential for dendritic cell maturation. (**a**) Immunofluorescence staining and quantification of cholera toxin B subunit (CTxB) as a marker for lipid nanodomains on the cell surface of conventional dendritic cells (cDCs) (unfed at steady-state) and mregDCs (DCs co-cultured with apoptotic cells) generated *in vitro*. (**b**) Flow cytometric staining for CTxB on the cell surface of CCR7^NEG^ and CCR7^POS^ cDCs *ex vivo* from lung tumors. Immunofluorescence staining for CTxB on the cell surface of (**c**) 1) cDCs, 2) cDCs supplemented with mevalonate, 3) mregDC, 4) mregDCs treated with the NPC1 inhibitor U18666A, 5) mregDCs treated with simvastatin, and 6) mregDCs treated with the LXR agonist GW3965, and on the cell surface of (**d**) (top) mregDCs treated with β-cyclodextrin and (bottom) mregDCs treated with sphingomyelinase. (**e**) Fluorescence quantification of CTxB staining on the groups shown in (**c**) and (**d**). (**f**) D4-GFP and CTxB quantification on wild-type and *Npc1*^+/-^ mregDCs by flow cytometry. Flow cytometric measurement of cDC maturation markers (i.e., CCR7, IFN-ɣR, CD40, CD80, CD86, PD-L1, PD-L2, MHC-I, MHC-II) on mregDCs either treated with (**g**) β-cyclodextrin (depletion of cholesterol from the plasma membrane) or (**h**) sphingomyelinase (selective depletion of cholesterol from lipid nanodomains). (**i**) Flow cytometric measurement of the cholesterol-independent receptor CD71 on mregDCs either treated with (left) β-cyclodextrin or (right) sphingomyelinase. Across all panels, data represent mean ± SEM. *P*-values computed by either unpaired *t-*test or ordinary one-way ANOVA *t*-test, as appropriate.

To identify a potential cDC-specific therapeutic target, we next sought to find candidate regulators of cholesterol transport in mregDCs. Given that efferocytosis drives the mregDC program and cholesterol is implicated in the underlying subcellular changes, we hypothesized that a efferocytosis-associated cell surface sensor may be responsible for the control of cholesterol mobilization in cDCs. Leveraging the integrated scRNAseq analysis, a survey of such receptors showed that, unlike most phagocytic receptors that are downregulated following antigen uptake, the expression of the receptor tyrosine kinase AXL was uniquely maintained on mregDCs (**Fig. 3a**). We hypothesized that this particular receptor may be involved in the engulfment and sensing of debris that subsequently modulates the intracellular dynamic of cholesterol in cDCs. By performing scRNAseq on MHC-II^+^ and CD11c^+^ cells from the tumor-bearing lungs of wild-type (*Axl*^+/+^, WT) and *Axl*-knockout (*Axl*^-/-^, KO) mice ^25^, we found that *Axl*-deficient mregDCs upregulated the *Npc1* gene and others encoding enzymes involved in *de novo* cholesterol synthesis, suggesting that AXL constitutively suppresses cholesterol mobilization upon uptake of cell debris (**Fig. 3b**,**c**). To confirm this, we used the D4-GFP probe to measure membranous free cholesterol on the surface *of Axl*^+/+^ *and Axl*^-/-^ mregDCs and observed a significant increase in signal from *Axl*-deficient mregDCs, compared to wild-type mregDCs (**Fig. 3d**, **Extended Data Fig. 3a**). These results were reproduced with direct fluorometric methods, as well (**Fig. 3e**). To determine whether the accumulating cholesterol observed in *Axl*-deficient mregDCs was derived from the phagocytosed apoptotic debris, we used the D4-GFP probe to first label the apoptotic bodies, prior to feeding them to cDCs. Here, too, significantly greater levels of GFP were measured in *Axl*-deficient mregDCs, compared to *Axl*-proficient mregDCs (**Extended Data Fig. 3b**), suggesting that in addition to newly synthesized cholesterol, a notable proportion of the mobilized cholesterol in mregDCs was derived from captured cargo and that AXL predominantly acts to limit the transport of extracellularly-derived cholesterol to the plasma membrane. By measuring the expression of maturation markers in *Axl*^+/+^ and *Axl*^-/-^ mregDCs, we observed that our findings on AXL and cholesterol translates to changes in maturation, as the expression of these markers was significantly elevated, in the absence of AXL (**Extended Data Fig. 3c-g**).

**Fig. 3.**
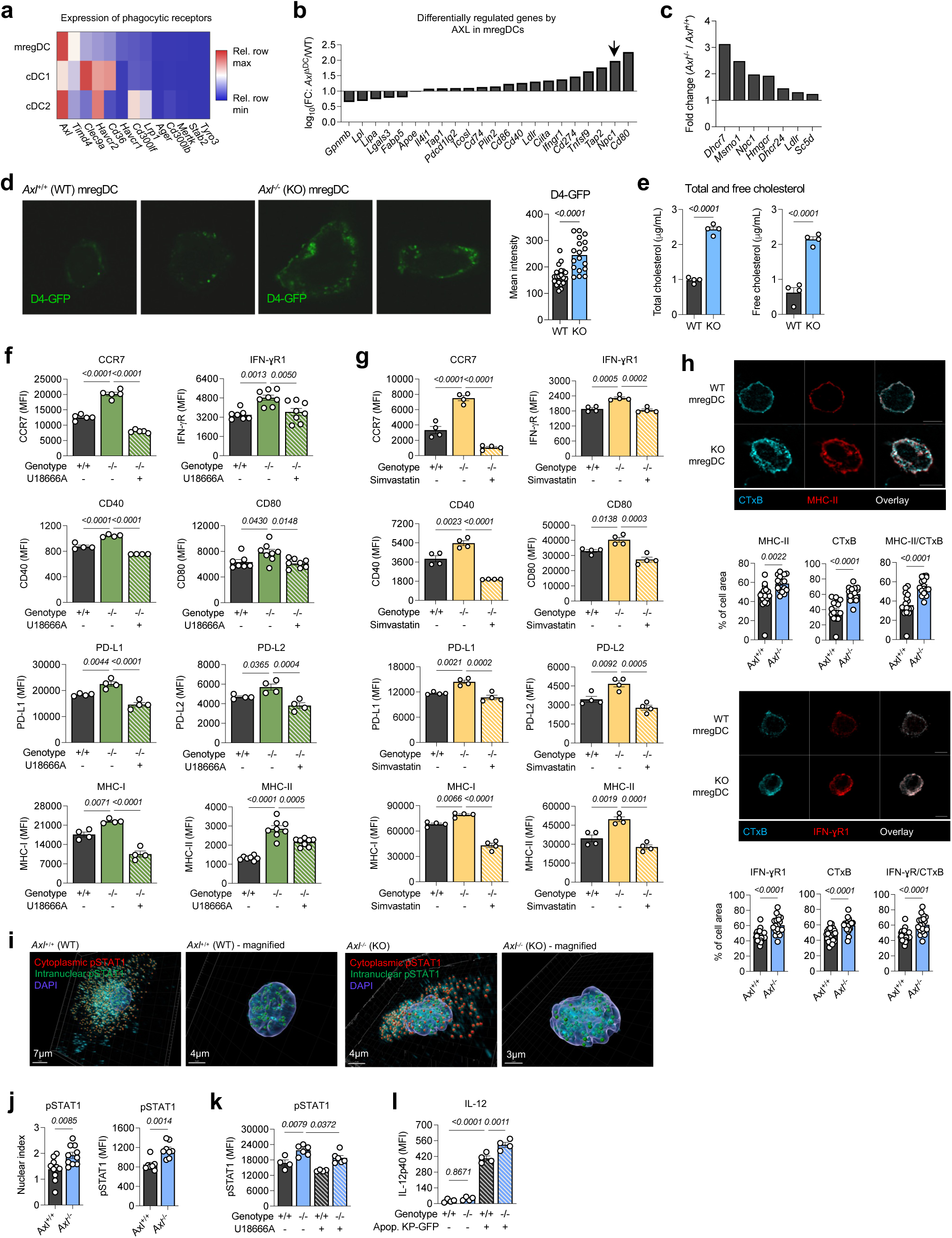
AXL limits cholesterol mobilization, lipid nanodomain assembly, and immunogenic receptor function. **(a)** Heatmap showing gene expression of targets encoding a select list of phagocytic cell sensors, computed relative to the row mean, across mregDC, cDC1, and cDC2 from the integrated analysis in Figure 1. (**b**) Differentially expressed genes between *Axl*^+/+^ (WT) and *Axl*^-/-^ (KO) mregDCs, as determined from mRNA profiling of myeloid cells in the lungs of WT and KO mice. (**c**) Differentially expressed genes associated with cholesterol neosynthesis, identified by topological analyses involving direct computational KEGG pathway mapping of differentially expressed genes between WT and KO mregDCs. (**d**) Immunofluorescence staining of D4-GFP-labelled WT and KO mregDCs and quantification of GFP intensity. (**e**) Fluorometric quantification of total and free cholesterol in WT and KO mregDCs. (**f**) Flow cytometric measurement of cDC maturation markers (i.e., CCR7, IFN-ɣR, CD40, CD80, MHC-I, MHC-II, PD-L1, and PD-L2) on WT and KO mregDCs either left untreated or treated with the NPC1 inhibitor U18666A. (**g**) Flow cytometric measurement of cDC maturation markers (i.e., CCR7, IFN-ɣR, CD40, CD80, CD86, PD-L1, PD-L2, MHC-I, and MHC-II) on WT and KO splenic mregDCs either left untreated or treated with simvastatin. (**h**) (Left) (Top) Localization and (bottom) quantification of MHC-II (red) and lipid nanodomains (CTxB, blue) on non-permeabilized WT and KO bone marrow-derived mregDCs; (right) (top) localization and (bottom) quantification of IFN-ɣR (red) and lipid nanodomains (CTxB, blue) on non-permeabilized WT and KO bone marrow-derived mregDCs. **(b)** (h) Confocal staining for intracellular phosphorylated STAT1 (pSTAT1) in WT and KO bone marrow-derived mregDCs. (h) Confocal imaging measurement (left) and flow cytometric measurement (right) of intracellular pSTAT1 and nuclear-to-cytosolic fluorescence ratio of pSTAT1 in WT and KO mregDCs. (**k**) Flow cytometric measurement of pSTAT1 in WT and KO splenic mregDCs either left untreated or treated with U18666A. (**l**) Flow cytometric measurement of intracellular IL-12p40 production by WT and KO splenic mregDCs that were stimulated with exogenous IFN-ɣ. Across all panels, data represent mean ± SEM. *P*-values computed by either unpaired *t-*test or ordinary one-way ANOVA *t*-test, as appropriate.

Given these results and the strong dependence of lipid nanodomain assembly on mobilizing cholesterol (**Fig. 2i**), we postulated that both the intracellular trafficking of cholesterol by NPC1 and its *de novo* synthesis are linked to the enhanced expression of maturation markers on *Axl*-deficient mregDCs. To test this, we treated *Axl*-deficient cDCs with either U18666A or simvastatin prior to feeding them with GFP^+^ apoptotic bodies and compared them with controls. Strikingly, we found that both NPC1-inhibited and cholesterol synthesis-impaired, *Axl-*deficient mregDCs exhibited abrogated expression of all maturation markers – including the immuno-stimulatory, -inhibitory molecules, and MHC-I and -II – relative to control *Axl*-deficient mregDCs (**Fig. 3f, g**). These results supported the hypothesis that there is a strong regulatory link between AXL and the mobilization of cholesterol through both its transport and *de novo* synthesis.

As modulation of lipidic species – including cholesterol and its oxidized derivatives – has been shown to perturb the localization and function of immunogenic receptors like MHC-II and IFN-ɣR ^1,4,9,18,36^, we hypothesized that the enhanced expression of maturation markers in the absence of AXL could be mediated by the improved stabilization of these immunogenic receptors *via* the cholesterol-based lipid nanodomains. We observed an increase in CTxB-occupied surface area on *Axl*-deficient mregDCs (**Fig. 3h**). Furthermore, these cholesterol-based lipid nanodomains ^15^ on the plasma membrane were specifically enriched with either MHC-II or IFN-ɣR1, compared to *Axl*-proficient cells (**Fig. 3h**).

We then demonstrated the functional implications of these findings by measuring phosphorylated STAT1 (pSTAT1) in *Axl*-deficient mregDCs, compared to *Axl*-proficient mregDCs, as a surrogate of IFN-ɣ signaling through its cognate receptor ^33^. Indeed, both flow cytometric and confocal imaging of cytosolic versus nuclear/translocated pSTAT1 showed enhanced signaling in mregDCs, in the absence of AXL (**Fig. 3i, j** and **Supplementary Videos 1,2**). Accordingly, the elevated pSTAT1 signal in the absence of AXL was compromised by U18666A, thus confirming the dependence of proper receptor localization and improved stabilization *via* lipid nanodomains on the AXL-and NPC1-regulated mobilization of cholesterol (**Fig. 3k**). Moreover, to determine whether this enhanced signaling response elicits greater production of immunostimulatory cytokines, we measured the T cell-activating cytokine IL-12 and found significantly greater production by *Axl*-deficient mregDCs (**Fig. 3l**), supporting the contribution of NPC1-dependent regulation by AXL to the immunogenicity of mregDCs.

We also assessed the antigen-presenting capacity of these cells. We found that the ability for mregDCs to activate OT-I or OT-II cells was significantly compromised in the presence of the different molecular inhibitors that reduced either the intracellular cholesterol reservoir or inhibited transport within cDCs (**Fig. 4a**). These results were confirmed using cDCs with a genetic deficiency for NPC1 (**Fig. 4b**) and cDCs without organized lipid nanodomains (**Fig. 4c**). In the absence of AXL, however, mregDCs exhibited elevated activation of CD8 and CD4 T cells. This was dependent on both cholesterol synthesis and transport, validating the functional relevance of an AXL-centric regulation of the cholesterol-based assembly of lipid nanodomains in cDC maturation and function (**Fig. 4d**). To establish the relevance of cholesterol in cDC maturation *in* vivo, we leveraged the maturation of cDCs in wild-type (WT) and *Npc1*^+/-^ mice that were challenged intravenously with 4T1 breast cancer cells to model a setting of excess cell debris in the lungs. Consistent with our proposed role for cholesterol in cDC maturation, we found that NPC1-deficient mregDCs in lung metastases of *Npc1*^+/-^ mice exhibited reduced expression of maturation markers, compared to their WT counterparts (**Extended Data Fig. 4a-d**). This suggested that impeding cholesterol mobilization in cDCs results in impaired maturation *in vivo*. We found that the regulatory function of AXL could also be appreciated *in vivo*. By first examining T cells in the lungs and mediastinal lymph nodes of naïve mice, we observed no significant differences among regulatory T cells (Tregs) or activated T cells (**Extended Data Fig. 5a**). However, a significantly greater proportion of CD4 and CD8 T cells in the lymph nodes exhibited activation potential (IFN-ɣ^+^ TNF-⍺^+^), indicative of a potential homeostatic role for AXL in promoting tolerance at the steady-state (**Extended Data Fig. 5b,c**).

**Fig. 4.**
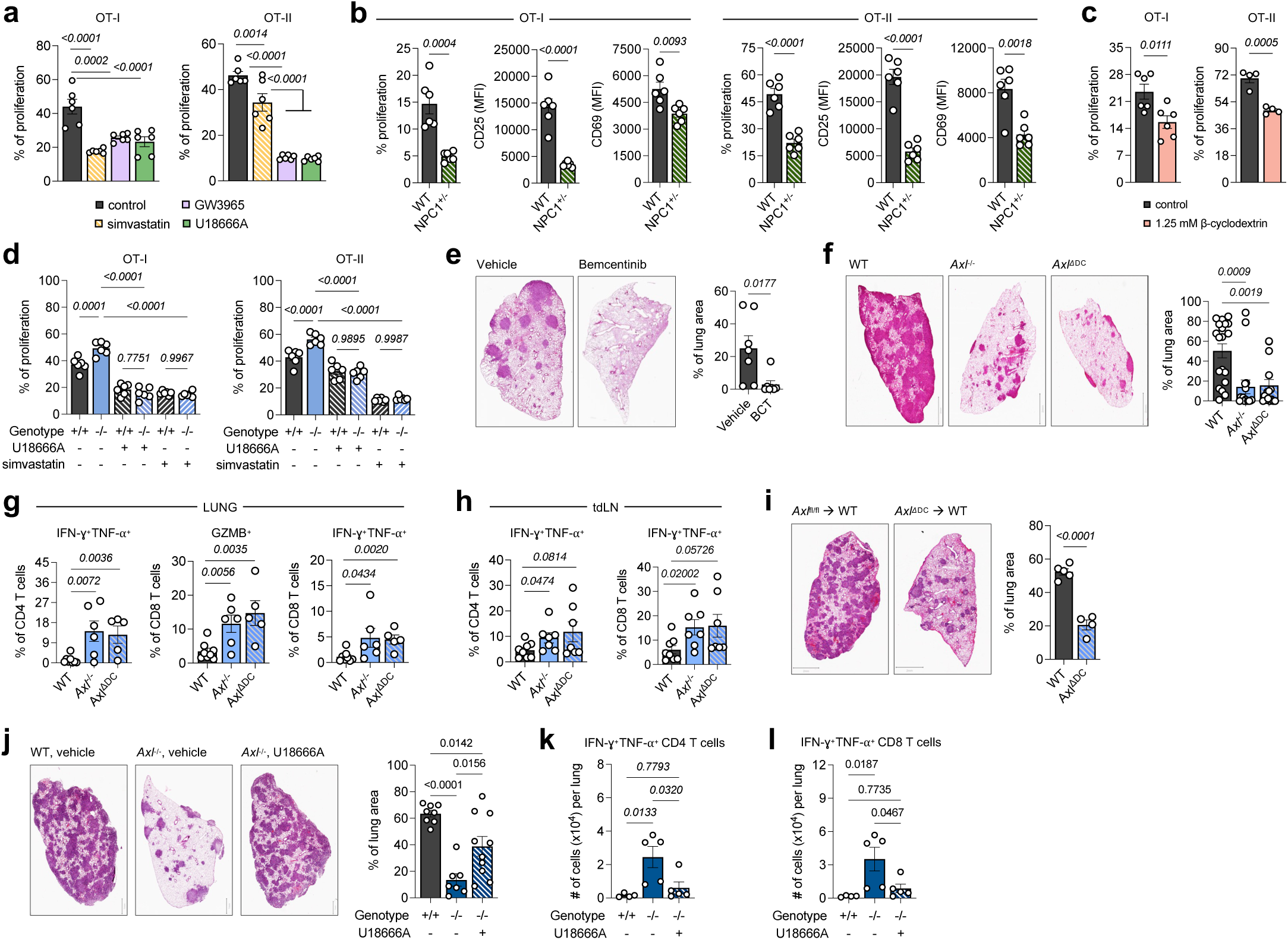
The AXL-NPC1 axis in cDC maturation controls anti-tumor T cell immunity. (**a**) *In vitro* antigen presentation efficacy of mregDCs either with control (PBS), simvastatin, GW3965 (the LXR agonist), U18666A (the NPC1 inhibitor), and fatostatin, as measured by the frequency of induced proliferation amongst (left) OT-I or (right) OT-II cells. Proliferation and activation of OT-I and OT-II cells, either (**b**) following co-culture with wild-type (WT) and *Npc1* haploinsufficient (*Npc1*^+/-^) mregDCs or **(c**) following co-culture with WT mregDCs treated with control or methyl-β-cyclodextrin. (**d**) WT and *Axl*^-/-^ (KO) mregDCs were generated by feeding cDCs of both genotypes with apoptotic tumor cells expressing OVA. *In vitro* Ag presentation efficacy in the presence of control (PBS), U18666A, or simvastatin was measured by the frequency of induced proliferation amongst (left) OT-I or (right) OT-II cells. (**e**) Tumor burden in the lungs of WT mice either treated with a vehicle or bemcentinib (AXL inhibitor), and (**f**) tumor burden in the lungs of WT, KO, and *Zbtb46*^Cre^-*Axl*^fl/fl^ (*Axl*^ΔDC^) mice at 21 days post-inoculation. (**g**) Frequency of activated and cytotoxic CD4 and CD8 T cells in lung tumors of WT, KO, and *Axl*^ΔDC^ mice. (**h**) Frequency of activated and cytotoxic CD4 and CD8 T cells in the tumor-draining lymph nodes (tdLNs) or WT, KO, and *Axl*^ΔDC^ mice. (**i**) Tumor burden in the lungs of recipient mice of *Axl*^fl/fl^ and *Axl*^ΔDC^ donor bone marrow. (**j**) Tumor burden in the lungs of WT, KO, and KO mice treated with U18666A at 17 days post-inoculation. Frequency of (**k**) activated CD4 and (**l**) cytotoxic CD8 T cells in lung tumors of mice shown in (**j**). Across all panels, data represent mean ± SEM. *P*-values computed by ordinary one-way ANOVA *t*-test.

As currently available modalities for perturbing cholesterol mobilization specifically in cDCs *in vivo* are limited, we reasoned that targeting AXL, which is more restricted in expression according to our query of the ImmGen Consortium database, is a more practical alternative to probing the AXL-cholesterol axis *in vivo*. So, to determine whether inhibiting AXL enhances mregDC immunogenicity and function *in vivo* in an otherwise immunosuppressive disease condition, we tested the pharmacological inhibition of AXL in the solid tumor context using our pre-clinical model of lung adenocarcinoma and the drug bemcentinib, a selective inhibitor for AXL, as AXL-targeting therapy has begun prevalent testing in clinical trials, many of which focus on lung cancer (NCT04681311, NCT01639508, NCT02988817 and NCT02729298). Briefly, *Kras*^G12D/+^*p53*^-/-^ (KP) lung epithelial cells were intravenously injected into *Axl*^+/+^ mice and were either treated with a control or bemcentinib, starting at 14 days post-inoculation with tumor cells. Histological exam of lung sections from the tumor-bearing lungs of control and bemcentinib-treated mice revealed significant elimination of tumor lesions due to AXL inhibition (**Fig. 4e**). However, as KP cells express AXL (**Extended Data Fig. 6a**) ^29^ and AXL has been shown to modulate tumor cell invasiveness ^6,10^, it remained possible for the therapeutic effect of AXL inhibition to have been conferred by a direct of bemcentinib on the tumor cells. To address this possibility, we assessed tumor growth in *Axl*^-/-^ mice and mice with a specific deletion of AXL in cDCs (*Zbtb46*^Cre^-*Axl*^fl/fl^ or *Axl*^ΔDC^) ^48^. We found that *Axl*^-/-^ mice and, more importantly, *Axl*^ΔDC^ mice showed significant reduction of tumor lesions in the lungs (**Fig. 4f**). *Ex vivo* staining for CTxB confirmed that *Axl*-deficient mregDCs indeed show enhanced assembly of lipid nanodomains on the cell surface, compared to proficient ones (**Extended Data Fig. 6b**); and accordingly, *Axl*-deficient mregDCs or mregDCs from mice treated with bemcentinib produced more IL-12, and significantly more CD8 and CD4 T cells in the tumor-bearing lungs and dLNs of *Axl*^-/-^ and *Axl*^ΔDC^ mice produced the effector cytokines IFN-ɣ and TNF-⍺ (**Extended Data Fig. 6c** and **Fig. 4g,h**). No differences in the frequency of NK cells in neither the lungs nor dLNs were documented (**Extended Data Fig. 6d**). We sought to determine whether this enhanced anti-tumor response was dependent on the robust T cell activity. Depletion of either CD8 or CD4 T cells abrogated the therapeutic effect of AXL deficiency, with a larger effect observed in the absence of CD4 T cells (**Extended Data Fig. 6e-h**), suggesting a particularly important role for CD4 T cells in “responding” to the enhanced immunogenicity of *Axl*-deficient mregDCs and coordinating a more potent anti-tumor immune response with CD8 T cells.

As AXL is also expressed on the endothelium and the *Zbtb46*^Cre^ is active in both cDC and endothelial cells ^62^, we sought to transplant bone marrow from *Axl*^ΔDC^ mice into wild-type mice to restrict AXL deficiency to hematopoietic cells (i.e. cDCs only) and assessed tumor burden in these mice, relative to control recipient mice that were given *Axl*^fl/fl^ marrow. We still observed a significant reduction in tumor burden in mice with *Axl*^ΔDC^ marrow (**Fig. 4i**), fully demonstrating the specific importance of AXL on cDCs in the enhanced anti-tumor immune response. Finally, to determine the *in vivo* relevance of a link between AXL and cholesterol in cDCs, we treated AXL-deficient mice with the U18666A compound. Quite strikingly, we observed that NPC1 inhibition in AXL-deficient mice significantly abrogated the therapeutic effect of AXL deficiency on tumor growth (**Fig. 4j**). This was accompanied by a complete impairment of both cytotoxic CD4 and CD8 T cell activity (**Fig. 4k,l**), which suggested to us that there is a robust axis between AXL and cholesterol *in vivo* that regulates mregDC immunogenicity.

Finally, we sought to establish the relevance of our *in vitro* and *in vivo* observations to human cDC biology using an *in vitro* culture system for human cDCs from CD34^+^ cord blood cells ^63^. We first performed bulk RNA-sequencing of sorted human cDC1 and mregDCs from a culture of cDCs that were fed apoptotic bodies. Leveraging our transcriptomic profiling of murine cDCs and mregDCs, we established that human mregDCs are homologous to the murine mregDCs (**Fig. 5a**), indicative of the utility of our *in vitro* system for generating human cDCs. Specifically, we could find that genes encoding enzymes involved in *de novo* cholesterol synthesis, classical markers of maturation, and AXL are all up-regulated in human mregDCs, compared to cDCs (**Fig. 5b**). Based on this data, we used the *in vitro* generation of human cDCs to test the effect of NPC1 inhibition on maturation. Strikingly, treating human mregDCs with the U18666A compound significantly reduced the expression of migratory (CCR7), co-stimulatory (CD80, CD83), and co-inhibitory (PD-L1, PD-L2) markers of maturation (**Fig. 5c**). Importantly, we also found that NPC1 inhibition reduced the ability for human cDCs to cross-present NY-ESO-1^+^ apoptotic cargo to NY-ESO-1-specific CD8 T cells (**Fig. 5d**), further demonstrating the functional relevance of cholesterol mobilization in human cDCs to the regulation of their immunogenic response.

**Fig. 5.**
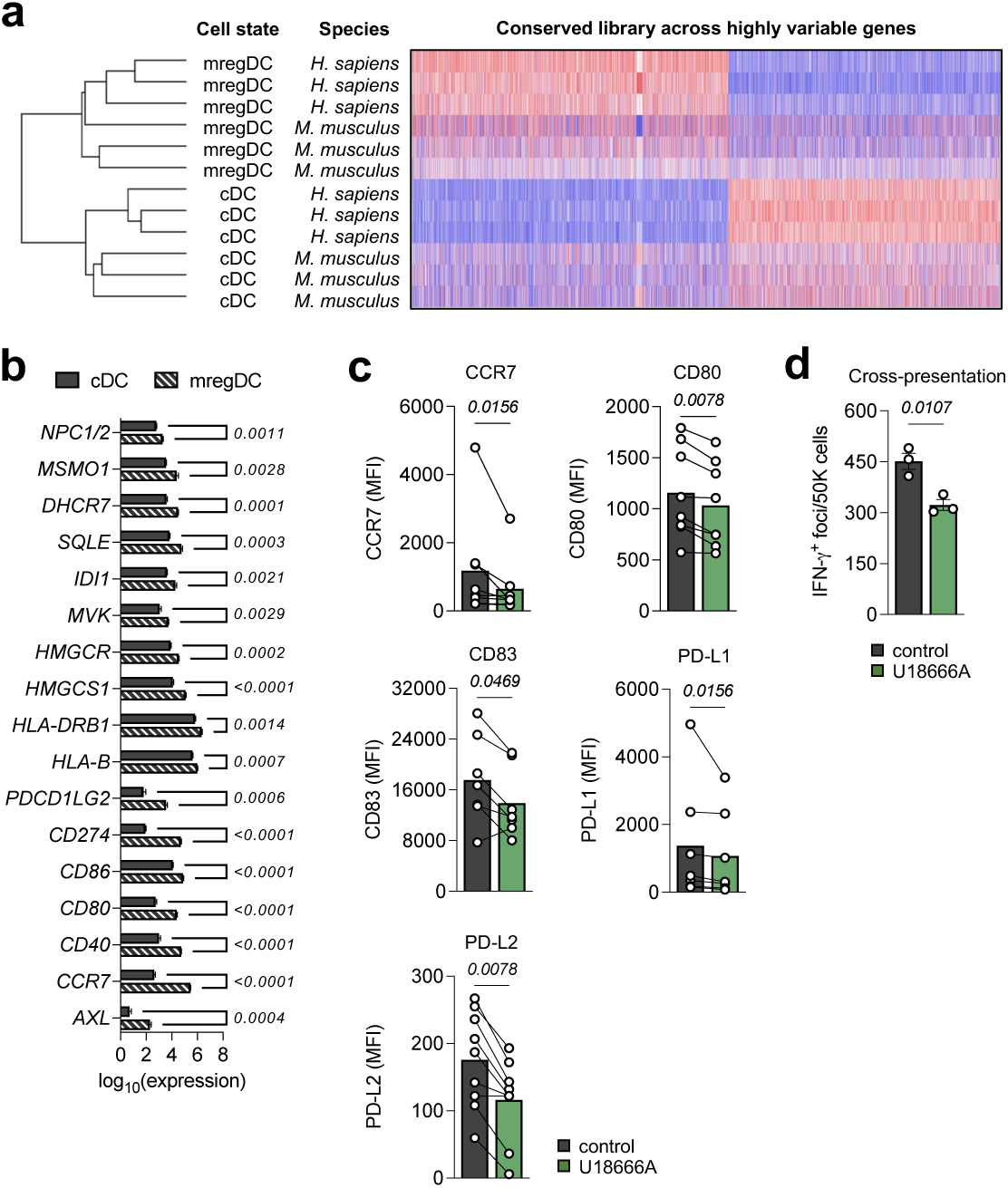
The effects of cholesterol mobilization on cDC maturation is conserved in humans. (**a**) Hierarchical clustering of sorted murine (*M. musculus*) and human (*H. sapiens*) cDCs and mregDCs following bulk RNA-sequencing, illustrating homology of transcriptomes. (**b**) mRNA expression of cDC maturation and *de novo* cholesterol synthesis genes by human cDCs and mregDCs. (**c**) Cell surface protein expression of cDC maturation markers by human mregDCs either treated with control (PBS) or U18666A. (**d**) *In vitro* antigen presentation efficacy of human mregDCs either treated with control (PBS) or U18666A, as measured by IFN-ɣ production by human CD8 T cells. Briefly, human mregDCs – either in the presence of control (PBS) or U18666A – were fed apoptotic K562 cells overexpressing the full-length NY-ESO1-p2A-eGFP and were then co-cultured with autologous NY-ESO1-specific CD8 T cells at a 1:1 ratio for 24 hours. Across all panels, data represent mean ± SEM. For panel (**b**) *p*-values computed by multiple unpaired *t*-tests. For panel (**c**), *p*-values computed by Wilcoxon matched-pairs signed rank test. For panel (**d**), *p*-value computed by unpaired *t*-test.

Taken altogether, our findings identify a new metabolic axis that governs the expression of maturation markers following the uptake of apoptotic cargo by cDCs. Transformation to the mregDC state is mediated by the mobilization of newly acquired and synthesized cholesterol that is trafficked and organized as lipid nanodomains on the plasma membrane of mregDCs. This response is critically dependent on NPC1, and the receptor tyrosine kinase AXL plays an important role in regulating it and the trafficking of cholesterol from *de novo* synthesis. AXL had previously been shown to induce the suppressors of cytokine signaling ^44^, and accordingly, enhanced expression of both *SOCS2* and *SOCS3* belong to the mregDC gene program. Here, we show that AXL suppresses NPC1 and *de novo* cholesterol synthesis to deter the assembly of lipid nanodomains and the expression of CCR7, MHC molecules and the IFN-ɣ receptor, among others, thus hampering the immunogenic potential of mregDCs. We showed that blocking AXL signaling releases a break on (*a*) the cDC response to IFN-ɣ, (*b*) their subsequent production of the immunostimulatory cytokine IL-12, and (*c*) the antigen presentation capacity to T cells. In sum, we have expanded the paradigm of cDC maturation by specifying the subcellular role for cholesterol dynamics, identifying lipid nanodomains as structural and functional features of mregDCs, and revealing a constitutive regulatory role for AXL in the expression and function of markers of cDC maturation. In combination with data generated with human cDCs, the results also proffer translational evidence that leveraging AXL as a therapeutic cDC target for enhancing mregDC immunogenicity and eliciting a remarkably strong anti-tumor immune response. Importantly, given the assessment of AXL in the clinic for its potential effect on tumors expressing AXL to modulate invasiveness and previous preclinical studies describing AXL as a macrophage target ^13,20,21,38,54^, our data not only present a novel role for AXL as a dendritic cell modulator of maturation, but also provide additional rationale for prioritizing the inhibition of AXL in lung cancer patients, given its dual effect on cancer cells and anti-tumor immunity.

## Materials and Methods

### Mice

*Axl*^fl/fl^ mice (a gift from Dr. Carla Rothlin, Yale University) were crossed with *Zbtb46*^Cre^ mice (JAX #028538) to generate *Axl*^ΔDC^ mice. *Axl*^fl/fl^ mice littermates were used as controls. *Axl*^-/-^ mice (complete KO) were purchased from Jackson Laboratories (JAX #005777) and bred in our facility. Transgenic OT-I TCR mice (JAX #003831), OT-II TCR mice (Taconic #11490), and *Npc1*^+/-^ mice (JAX #003092) were used for experiments after at least one week of housing in our facility. All mice were maintained at specific pathogen-free (SPF) health status in individually ventilated cages at 21-22°C and 39-50% humidity and were handled in compliance with national and institutional guidelines. Mice used in experiments were between 8 and 12 weeks old.

### *In vivo* model of lung cancer

The GFP-expressing *Kras*^G12D/+^ *p53*^-/-^ (KP) or *Kras*^G12D/+^ *p53*^-/-^ *Rosa26*^A3Bi^;*Rag1*^-/-^ (KPAR) cell lines were tested negative for mycoplasma and validated for its potential to generate lung tumors. Eight-week-old mice were inoculated intravenously with either 5 x 10^5^ KP-GFP or 1.5 x 10^4^ KPAR cells. Lungs and lymph nodes were analyzed on day 21. When indicated, tumor-bearing mice were treated with either vehicle or bemcentinib (BerGenBio) by oral gavage twice a day starting at day 14 until day 21; or injected intraperitoneally (i.p.) with 200 μg/mL anti-CD8α-depleting antibody (BioXcell, clone 2.43) or 200 μg/mL anti-CD4-depleting antibody (BioXcell, clone GK1.5) on days 10, 13, 16 and 19. For systemic treatment with the NPC1 inhibitor U18666A, 50 micrograms were given to each mouse intraperitoneally. To quantify lung tumors, paraffin-embedded lung lobes were stained with hematoxylin & eosin (H&E); the slides were scanned using an Olympus digital scanner and analyzed using QuPath software.

### Reagents and antibodies

Human: Anti-human CD141 (clone M80, BioLegend), anti-human CD45RA (clone HI100, BioLegend), anti-human CD1C (clone L161, BioLegend), anti-human CD123 (clone 6H6, BioLegend), anti-human CLEC9A (clone 8F9, BioLegend), anti-human CADM1 (clone 30, MBL International), anti-human CD206 (clone 15-2, BioLegend), anti-human CD80 (clone W17149D, BioLegend), anti-human CD83 (clone HB15e, BioLegend), anti-human CD86 (clone BU63, BioLegend), anti-human PD-L1 (clone 29E2A3, BioLegend), anti-human PD-L2 (clone MIH18, BioLegend), anti-human HLA-A,B,C (clone W6/32, BioLegend), anti-human CCR7 (clone 3D12, Invitrogen), streptavidin (BioLegend). Anti-human IFN-ɣ (biotin) (clone 7-B6-1, mabtech) and (unconjugated) (clone 1-D1K, mabtech), streptavidin-AP conjugate (Roche), SIGMAFAST BCIP/NBT (Sigma-Aldrich). Mouse: anti-CD19 PerCP (Invitrogen), anti-F4/80 PerCP (Invitrogen), anti-CD11c APC-Cy7 (Invitrogen), anti-IA/IE eFluor450 (BioLegend), anti-CD11b AlexaFluor700 (Invitrogen), anti-CD8α BV785 (BioLegend), anti-F4/80 PE-Cy7 (BioLegend), anti-GR1 APC (eBioscience), anti-B220 APC (BioLegend), anti-CD3 APC (Invitrogen), anti-IA/IE AlexaFluor700 (Invitrogen), anti-CD8α eFluor450 (Invitrogen), anti-CD11b PE (eBioscience), anti-CD86 eFluor450 (BioLegend), anti-CD40 APC (Invitrogen), anti-CD80 Pacific Blue (BioLegend), anti-CD119 PE (Invitrogen), anti-CCR7 APC (BioLegend), anti-MHC-I PE (eBioscience), anti-CD74 APC (BioLegend), anti-IL12 PE (BioLegend), anti-pSTAT1 (BD Biosciences), CellTrace violet stain (Invitrogen), anti-cholera toxin B-subunit goat pAB (Sigma-Aldrich), anti-IA/IE biotin (Invitrogen), Hamster anti-CD119 (Santa Cruz Biotechnology), Streptavidin Alexa Fluor 555 (Invitrogen), Alexa Fluor 647 donkey anti-goat (Invitrogen), U18666A (Sigma-Aldrich), mevalonic acid (Sigma-Aldrich), D4-GFP (kindly provided by Dr. Christophe Lamaze), bemcentinib (kindly provided by BerGenBio), simvastatin (Cayman Chemical), methyl-β-cyclodextrin (Sigma), hydrolase sphingomyelinase (Sigma), GW3965 (Sigma).

### Dendritic cell purification and bone marrow-derived dendritic cell culture

Splenic and bone marrow-derived DC (BMDC) purification was performed, as described previously (Belabed et al., 2020). Briefly, spleens from wild-type or *Axl*^-/-^ mice were digested with 500 μg/mL Liberase (Roche Diagnostics) and 50 ng/mL recombinant DNase I (Roche Diagnostics) in PBS. The DCs were pre-enriched from splenocytes using a very-low-density gradient. Splenocytes were first resuspended in 4.2 mL of RPMI medium containing 5mM EDTA, 5% FBS and 1mL Optiprep (Sigma), then loaded between 3 mL PBS containing 5 mM EDTA, 5% FCS and 1 mL Optiprep (bottom layer) and 1.8 mL of PBS containing 5 mm EDTA and 5% FBS (top layer). After 20 min of centrifugation at 1800 rpm at room temperature, the low-density fraction was collected at the interface between the top and middle layers. For the generation of BMDCs, bone marrow cells were cultured for 8-10 days in medium supplemented with GM-CSF, 10% heat-inactivated FBS, 1% non-essential amino acids, 1mM sodium pyruvate, 100 U/mL penicillin and 100 U/mL streptomycin.

The cells were immuno-stained with the following antibodies: anti-CD19 PerCP, anti-F4/80 PerCP, anti-CD11c APC-Cy7, anti-IA/IE eFluor450 by flow cytometry.

### Efferocytosis assay

For induction of apoptosis, KP-GFP cells or B16-Ova-BFP were ultraviolet-irradiated 30 min and incubated overnight at 37°C with 5% CO_2_. When indicated, DCs were treated overnight with either 400 μM mevalonic acid, 50 µM simvastatin (Cayman Chemical), 0.625 mM methyl-β-cyclodextrin (Sigma), 0.3 unit/mL of hydrolase sphingomyelinase (Sigma), 40 µM GW3965 (Sigma), or 7.5 μg/mL of U18666A and equimolar volumes of DMSO as control. DCs were incubated with the apoptotic cells at a 1:3 DC:apoptotic cell ratio for 2h. Phagocytosis was assessed by flow cytometry by gating on GFP^+^ DCs or BFP^+^ DCs.

### *In vitro* cross-presentation assay

When indicated, DCs were treated overnight with either 400 μM mevalonic acid, 50 µM simvastatin (Cayman Chemical), 0.625 mM methyl-β-cyclodextrin (Sigma), 0.3 unit/mL of hydrolase sphingomyelinase (Sigma), 40 µM GW3965 (Sigma), or 7.5 μg/mL of U18666A and equimolar volumes of DMSO as control. DCs were incubated with apoptotic B16-OVA at a 1:2 (DC:apoptotic cell) ratio overnight. The cells were washed three times with PBS 0.5% BSA and co-cultured with OT-I cells (DC: OT-I ratio of 1:3) for 3 days to analyze OT-I cells by flow cytometry.

### OT-II presentation assay

For the *in vitro* OT-II presentation assay, when indicated, DCs were treated overnight with either 400 μM mevalonic acid, 50 µM simvastatin (Cayman Chemical), 0.625 mM methyl-β-cyclodextrin (Sigma), 0.3 unit/mL of hydrolase sphingomyelinase (Sigma), 40 µM GW3965 (Sigma), or 7.5 μg/mL of U18666A and equimolar volumes of DMSO as control. DCs were incubated with apoptotic B16-OVA at a 1:2 (DC:apoptotic cell) ratio overnight. The cells were washed three times with PBS 0.5% BSA and co-cultured with OT-II cells at a 1:3 (DC: OT-II) ratio for 3 days to analyze OT-I cells by flow cytometry.

### Sample preparation for single-cell RNA sequencing

Single-cell suspensions from lung tissues were obtained, as described above. For scRNAseq, these cells were suspended and stained in 100μL of multiplex hashing antibodies at 4°C for 20 minutes. Stained cells were washed three times in PBS+0.5% BSA to remove unbound antibodies. Washed cells were resuspended in 150μL of wash buffer and counted using a Nexcelom Cellometer Auto2000. Hashed samples were pooled in equal amounts of live cells. Volume was adjusted to achieve a target of 2×10^6^ cells/mL. Hashed samples were loaded onto 10X Genomics NextGen 5’ v1.1 assay, as per the manufacturer’s instructions, for a target cell recovery of 20,000 cells/lane. Libraries were constructed, as per the manufacturer’s instructions. During cDNA amplification, hashtag oligonucleotides (HTO) were enriched during cDNA amplification with the addition of 3 pmol of HTO Additive primer (5’GTGACTGGAGTTCAGACGTGTGCTC). This PCR product was isolated from the mRNA-derived cDNA via SPRISelect size selection, and libraries were made as per the New York Genome Center Hashing protocol. All libraries were quantified via Agilent 2100 hsDNA Bioanalyzer and KAPA library quantification kit (Roche, Cat. No. 0796014001.) Gene expression libraries were sequenced at a targeted depth of 25,000 reads per cells, and HTO libraries were sequenced at a targeted read depth of 1,000 reads per cell. All libraries were sequenced on the Illumina NovaSeq S2 100 cycle kit with run parameters set to 28×8×0×60 (R1xi7xi5xR2).

### scRNAseq analysis

After library demultiplexing, gene-expression libraries were aligned to the mm10 reference transcriptome and count matrices were generated using the default Cell Ranger 2.1 workflow, using the ‘raw’ matrix output. Where applicable, doublets were removed based on co-staining of distinct sample-barcoding (‘Hashing’) antibodies (maximum staining antibody counts/second-most staining antibody counts = less than 5). Following alignment, cell barcodes corresponding to cells that contained more than 500 UMIs were extracted. From among these, cells whose transcripts constituted more than 25% mitochondrial genes were filtered from downstream analyses. The R package *Seurat* was used to scale the data, transform via a log normalization method, adjust for batch correction, cluster cells based on shared nearest neighbors, and perform dimensionality reduction based on the first 15 principal components. Gene module analyses were performed, based on the identification of groups of highly correlated genes, according to the Pearson correlation matrix of the most variable genes. This was done using the R package *scDissector*. Differentially expressed genes were identified using the *FindMarkers* function in Seurat. Mean UMI expression values were imputed to determine log fold change differences between cell types to further the analysis of markers of interest.

### Confocal immunofluorescence

Bone marrow-derived DCs were plated on glass coverslips and fixed by incubation for 15 min on ice in 3.7% w/v paraformaldehyde, then quenched for 10 min with 50mM NH_4_Cl in PBS. The cells were incubated for 1 h with specific primary antibodies in PBS, washed twice, and incubated 1 h with the fluorescently labelled secondary antibody. Finally, the cells were mounted on slides in Prolong Gold antifade reagent in the presence of DAPI (Invitrogen Carlsbad, CA). Confocal microscopy was performed with a Zeiss LSM 780 system (Carl Zeiss) and a 20 or 63x objectives. Images were processed with the Zeiss LSM Image Browser (Carl Zeiss) and ImageJ software.

### Bone marrow transplantation

Recipient mice were irradiated with two doses of 5.5 Gy that were administered 6 hrs apart. Donor bone marrow cells (5×10^6^) were retro-orbitally transferred into irradiated recipient mice. A period of 7-8 weeks was granted to ensure engraftment. The recipients were supplemented with sulfamethoxazole / trimethoprim for three weeks. Reconstitution was assessed by flow cytometric analysis of inflammatory or Ly6C^HI^ monocytes and T cells in the lungs. Donor chimerism of at least 80% was confirmed for all mice.

### *In vitro culture of* human dendritic cells

Human cDCs were generated using hematopoietic stem cells derived from cord blood in a two-step culture system. CD34^+^ cells were expanded and differentiated with a combination of recombinant human cytokines in ⍺-MEM media supplemented with 10% FBS, as previously described ^63^. These cells were expanded for seven days with recombinant human Flt3L (25 ng/mL), SCF (2.5 ng/mL), TPO (5 ng/mL), and IL-7 (5 ng/mL). Expanded cells were differentiated in a feeder layer culture (OP9 + OP9 DL1) for 18 days with a combination of Flt3L (5 ng/mL), IL-7 (5 ng/mL), TPO (2.5 ng/mL), and GM-CSF (0.5 ng/mL). Cultures were replenished every seven days by removing half of the media and replacing it with 500 µL of fresh media with cytokines at twice their prescribed concentrations. DCs were identified based on the expression of CD141, CLEC9A, and CADM1 (cDC1), of CD1c and CD206 (cDC2), and of CD45RA and CD123 (pDC).

*In vitro* generated human DCs were treated with or without the NPC1 inhibitor for 24 hours and co-cultured with CellTrace Far Red labelled apoptotic GFP-expressing tumor cells for 16 hours. The cells were harvested and assessed for the expression of different maturation markers by flow cytometry.

To assess uptake of cell debris and antigen cross-presentation, MHC-I-deficient K562 cells were engineered to overexpress full length NY-ESO1-p2A-eGFP. These cells were then labelled with CellTrace Far Red dye and heat-killed by keeping them at 55 deg C for 30 min and frozen for future use. The K562 cells were thawed and co-cultured with *in vitro* generated cDC1s for 24 hrs. cDC1s were fed with dead K562-NY-ESO1-eGFP cells. Then, these cells were washed and co-cultured with NY-ESO1^157–165^ epitope-specific autologous CD8 T cells at a 1:1 ratio for an ELISPOT assay for 24 hours. ELISPOT plates were developed and quantified for IFN-ɣ spots using the Immunospot ELISPOT reader (Immunospot 7.0.15.1 software, Immunospot C.T.L. Cellular Technology).

### Bulk RNA-sequencing

Untreated cDC1s and fed cDC1s were sorted and total mRNA was extracted using the RNeasy Plus Micro Kit (QIAGEN). mRNA was processed for 50 base-pair paired-end sequencing using NovaSeq 6000.

### Data Availability

Single-cell RNA-sequencing data generated for this study will be uploaded to the Gene Expression Omnibus NCBI database and an accession code will be made available upon publication.

## Supplementary Information

**Supplementary Table 1.** Sub-modules defined by the identification of highly correlated variable genes across dendritic cells integrated from different single-cell RNA-sequencing datasets.

**Supplementary Table 2.** Metadata per cell barcode of integrated single-cell RNA-sequencing analysis, including cell subtype annotation and dataset of origin.

**Supplementary Table 3.** Active metabolic pathways inferred from gene set enrichment analysis of genes enriched in mregDCs.

**Supplementary Video 1.** Z-stack recording of pSTAT1 and DAPI staining of wild-type BMDCs fed with KP-GFP cells.

**Supplementary Video 2.** Z-stack recording of pSTAT1 and DAPI staining of *Axl* knock-out BMDCs fed with KP-GFP cells.

## Acknowledgements

We thank members of the Merad Laboratory for helpful discussions; Prerna Suri and Robert Samstein in the Department of Immunology and Immunotherapy for providing the 4T1 cells; Shilpa D. Kumar and the Advanced Microscopy and Bioimaging Core; and the Mount Sinai Flow Cytometry Core and Human Immune Monitoring Center for technical support.

## Author Information

### CONTRIBUTIONS

M.M. conceived the project. M.B. and M.M. designed the experiments. M.B., M.D.P., and M.M. wrote the manuscript.

M.B. performed experiments, with support from M.D.P., J.L.B., R.M., and C.M.W. K.J.R., S.B., and A.P. performed experiment on human cDCs, including RNA sequencing. M.D.P. performed computational analyses. C.M.B. generated the D4-GFP cholesterol probe. S.T.C. and N.M.L. contributed to the conceptual development of the project. D.P., S.G., C.V.R., C.M.B., and C.L. advised on study design. N.M.L. was supported by the Cancer Research Institute / Bristol Myers Squibb Irvington Postdoctoral Research Fellowship to Promote Racial Diversity (Award No. CRI3931). R.M. was supported by the 2021 AACR-AstraZeneca Immuno-oncology Research Fellowship, (Grant No. 21-40-12-MATT). K.J.R. was supported by a Fulbright Future Scholarship and the Mater Foundation. C.M.B. and C.L. were supported by institutional grants from the Curie Institute, INSERM, CNRS, and by grants from Agence Nationale de la Recherche (ANR NanoGammaR-17-CE15-0032) to C.L., from Ligue Nationale contre le Cancer and ARC to C.M.B. C.M.B. and C.L. are members of Labex CelTisPhyBio ANR-10-LBX-0038 and are part of the IDEX PSL ANR-10-IDEX-0001-02 18796.

---

**Extended Data Fig. 1.**
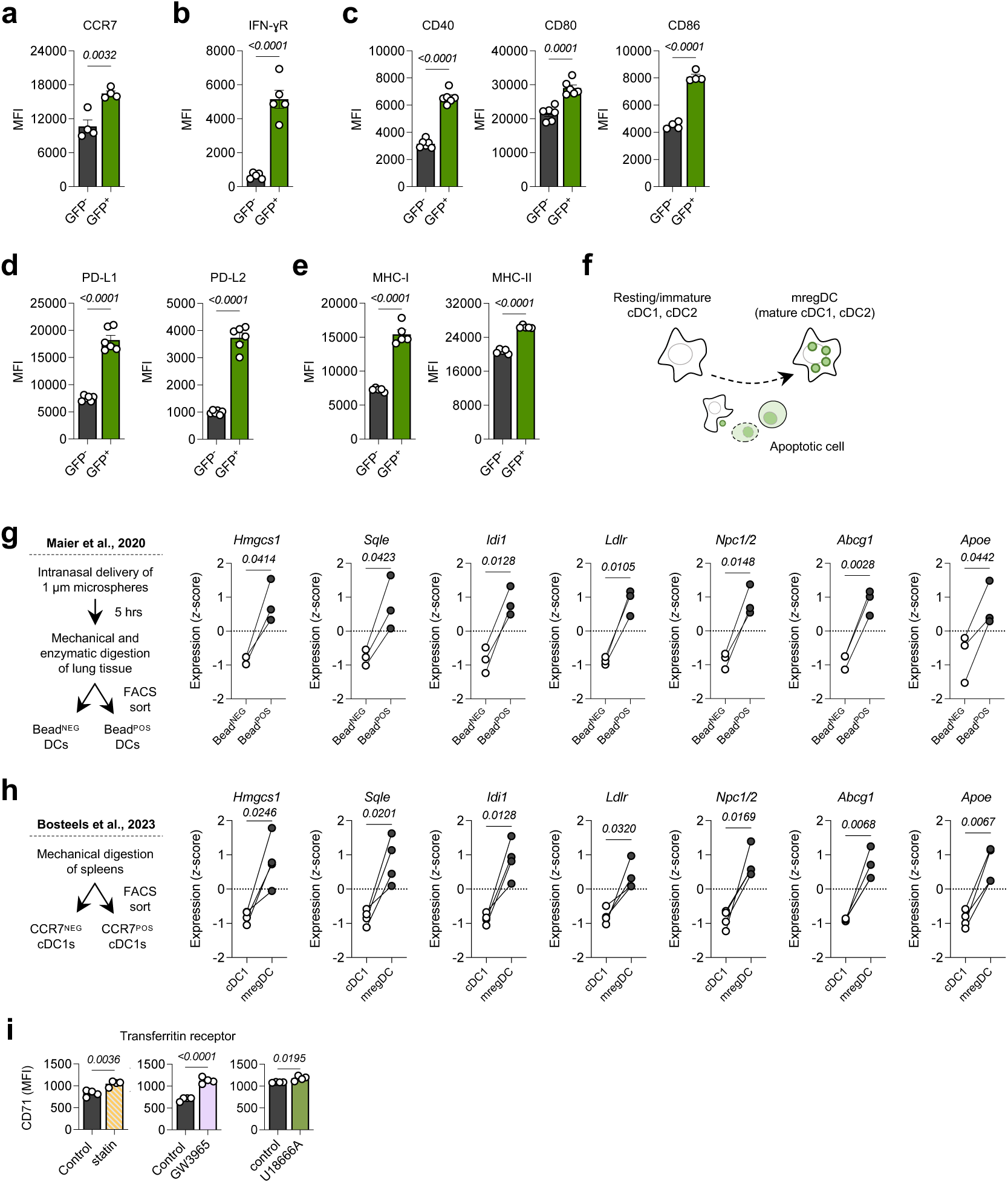
Uptake of cell debris triggers dendritic cell maturation and an mRNA signature of cholesterol mobilization. Flow cytometric measurement of (**a**) CCR7, (**b**) IFN-ɣ receptor, (**c**) CD40, CD80, and CD86, (**d**) PD-L1 and PD-L2, and (**e**) MHC class I and II on splenic cDCs either cultured alone or fed *in vitro* with apoptotic GFP-expressing tumor cells, where GFP functions as a surrogate antigen. (**f**) Graphical summary of the transformation of cDC maturation towards the mregDC state. (**g**) Gene expression of different genes encoding enzymes and transporters involved in cholesterol mobilization by FACS-sorted lung cDCs based on uptake status of microspheres administered intranasally *in vivo*. Data acquired from Maier et al., 2020. (**h**) Gene expression of different genes encoding enzymes and transporters involved in cholesterol mobilization by FACS-sorted splenic cDCs based on CCR7 expression. (**i**) Cell surface, cholesterol-independent expression of the transferritin receptor CD71 on mregDCs either treated with control (PBS), simvastatin, GW3965, or U18666A. Data for panels (**g**) and (**h**) were obtained from Maier et al., 2020 and Bosteels et al., 2023, respectively. Across all panels, data represent mean ± SEM. For panels (**a**)-(**d**), *p*-values computed by unpaired *t-*test. For panels (**e**), (**f**), *p*-values computed by paired *t*-test.

**Extended Data Fig. 2.**
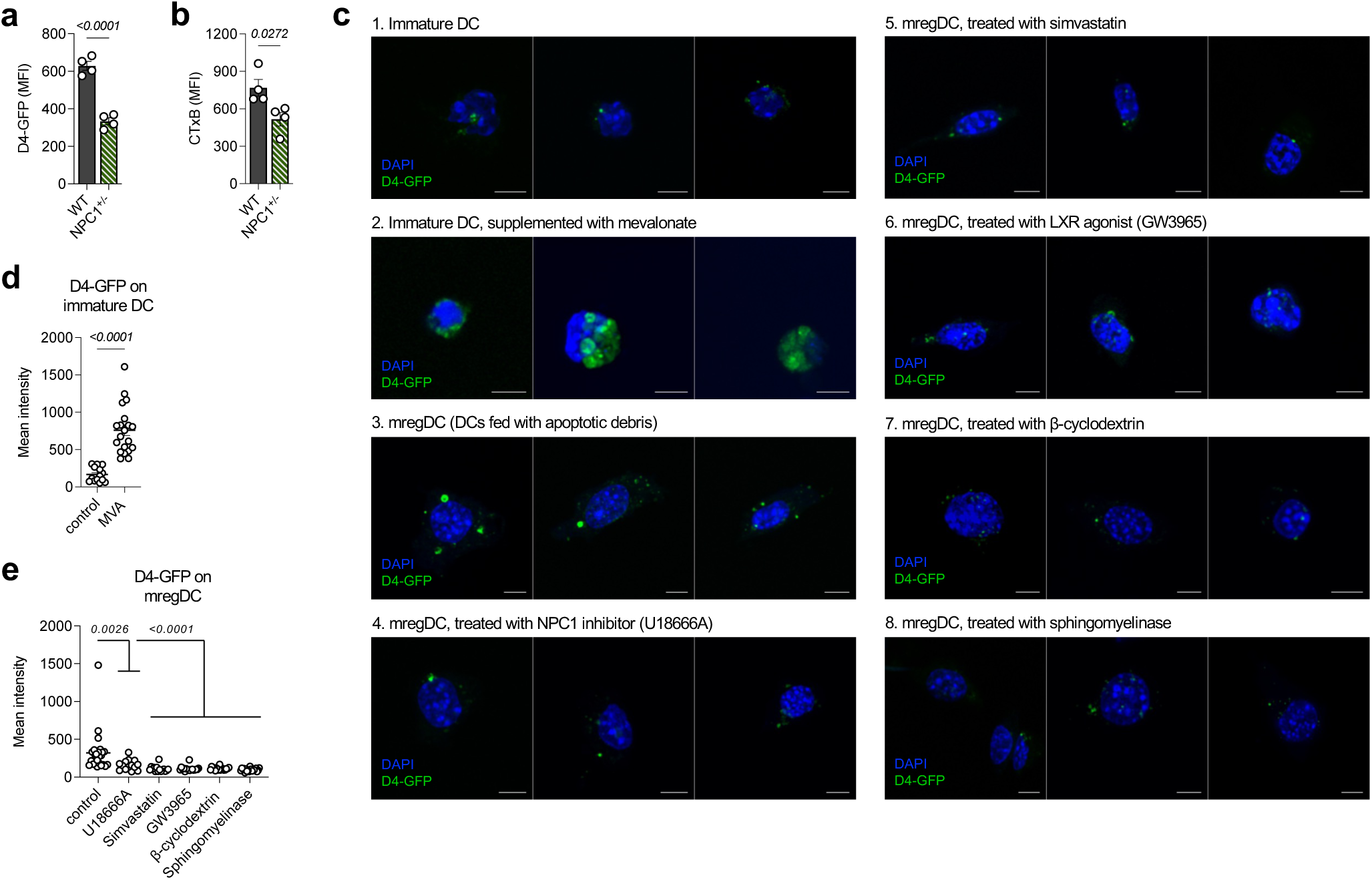
Quantification of cell surface cholesterol. Flow cytometric quantification of (**a**) D4-GFP and (**b**) CTxB staining of wild-type (WT) and *Npc1*^+/-^ mregDCs. (**c**) Immunofluorescence staining for free cholesterol on the cell surface of 1) cDCs, 2) cDCs supplemented with mevalonate, 3) mregDCs, 4) mregDCs treated with the NPC1 inhibitor U18666A, 5) mregDCs treated with simvastatin, and 6) mregDCs treated with the LXR agonist GW3965, 7) mregDCs treated with β-cyclodextrin, 8) mregDCs treated with sphingo-myelinase. (**d**) Fluorescence quantification of D4-GFP staining on cDCs and cDCs supplemented with mevalonate. (**e**) Fluorescence quantification of D4-GFP staining on groups 3-8 in (**c**). Across all panels, data represent mean ± SEM. *P*-values computed by unpaired *t-*test.

**Extended Data Fig. 3.**
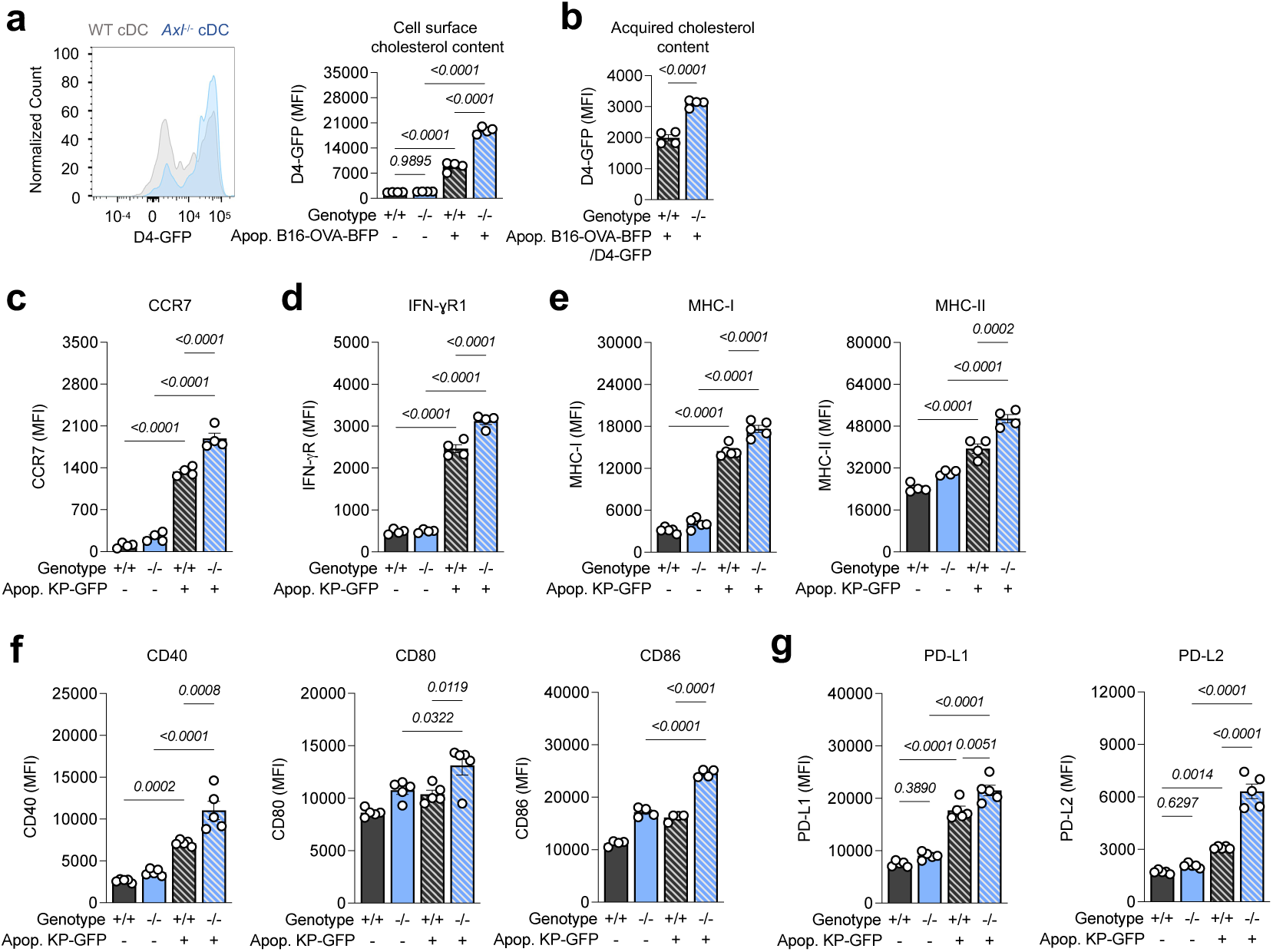
The phagocytic cell sensor AXL controls dendritic cell maturation. (**a**) Flow cytometric measurement of free cholesterol using the D4-GFP probe on WT and KO mregDCs. (**b**) Flow cytometric measurement of free cholesterol acquired extracellularly by mregDCs exposed to D4-GFP-stained apoptotic cells. (**c**)-(**g**) Flow cytometric measurement of cDC maturation markers (i.e., CCR7, IFN-ɣR, MHC-I, MHC-II, CD40, CD80, CD86, PD-L1, PD-L2). Across all panels, data represent mean ± SEM. *P*-values computed by ordinary one-way ANOVA *t*-test.

**Extended Data Fig. 4.**
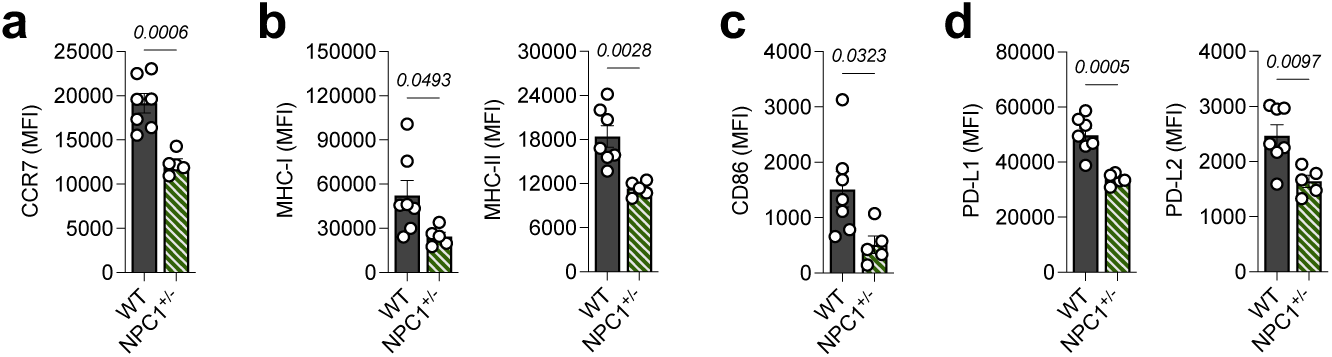
Dendritic cell maturation is dependent on NPC1 *in vivo*. Flow cytometric measurement of (**a**) CCR7, (**b**) MHC-I and MHC-II, (**c**) CD86, and (**d**) PD-L1 and PD-L2 on mregDCs isolated from tumor-bearing lungs in a 4T1 breast cancer metastasis model. Across all panels, data present mean ± SEM. All *p*-values computed by unpaired *t-*test.

**Extended Data Fig. 5.**
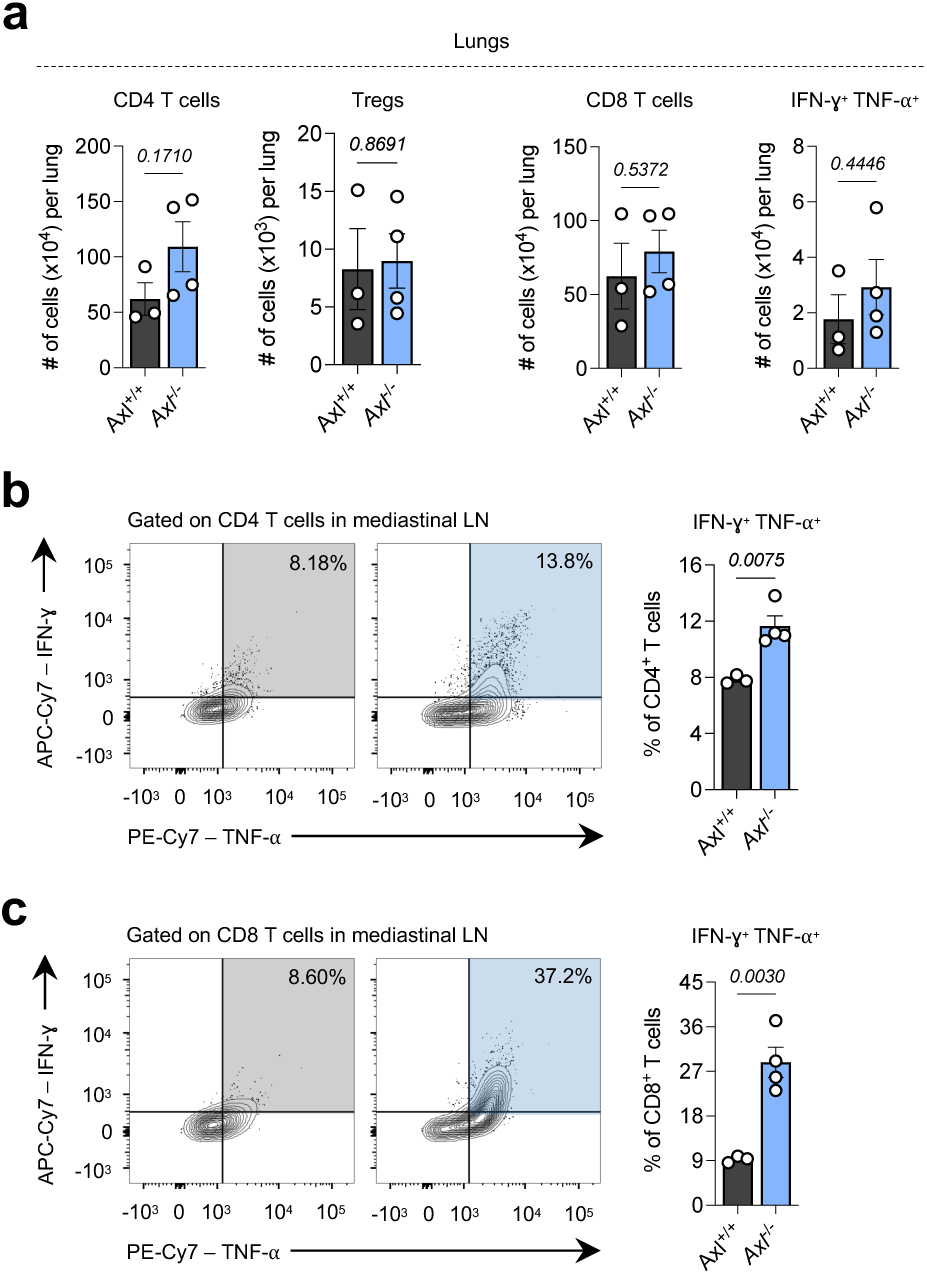
AXL can act as a dendritic cell checkpoint that promotes tolerance at steady-state. **(a)** Flow cytometric measurement of total CD4 T cells, regulatory T cells (Tregs), total CD8 T cells, and cytotoxic (IFN-ɣ^POS^ TNF-⍺^POS^) CD8 T cells in the lungs of naïve *Axl*^+/+^ (WT) and *Axl*^-/-^ (KO) mice. Flow cytometric measurement of (**b)** cytotoxic CD4 and (**c**) cytotoxic CD8 T cells in the mediastinal lymph nodes of naïve WT and KO mice. Across all panels, data represent mean ± SEM. All *p*-values computed by unpaired *t-*test.

**Extended Data Fig. 6.**
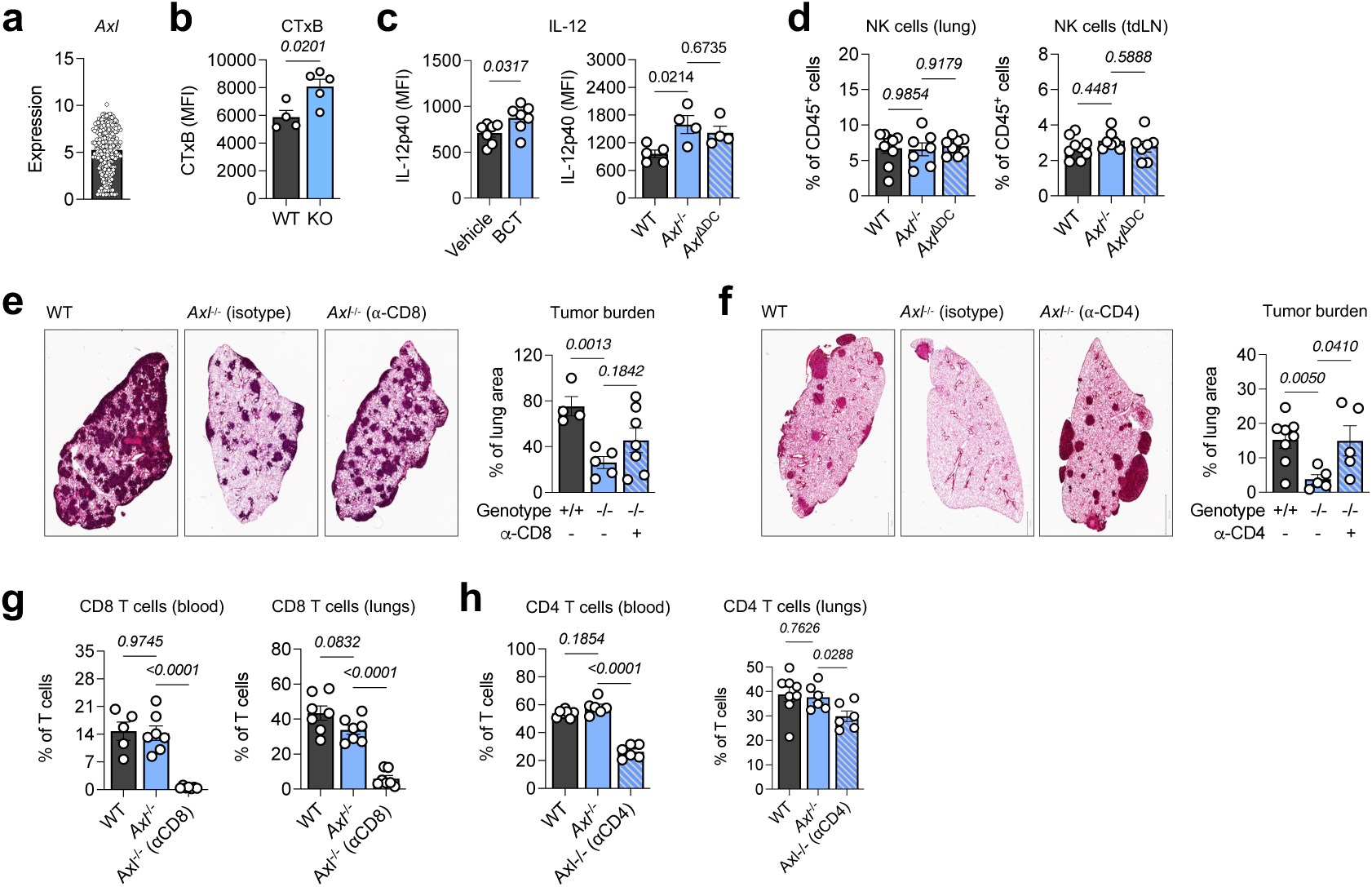
The therapeutic efficacy of the cDC checkpoint AXL is a T cell-dependent response. **(a)** mRNA expression of AXL by *Kras*^G12D/+^*Trp53*^-/-^ tumor cells, according to single-cell RNA-sequencing of tumor cells from the KP GEMM model of lung adenocarcinoma (Marjanovic et al., 2020). (**b**) *Ex vivo f*low cytometric staining of CTxB on *Axl*^+/+^ (WT) and *Axl*^-/-^ (KO) mregDCs from tumor-bearing lungs. (**c**) Flow cytometric measurement of intracellular IL-12p40 produced by mregDCs from the tumor-bearing lungs of (left) WT and BCT-treated mice or (right) WT, KO and *Zbtb46*^Cre^-*Axl*^fl/fl^ (*Axl*^ΔDC^) mice. (**d**) Frequency of NK cells in the (left) lungs and (right) tumor-draining lymph nodes of WT, KO, and *Axl*^ΔDC^ mice. (**e**) (Left) histology and (right) quantification of tumor burden in the lungs of WT, KO, and KO mice depleted of CD8 T cells at 21 days post-tumor cell inoculation. (**f**) (Left) histology and (right) quantification of tumor burden in the lungs of WT, KO, and KO mice depleted of CD4 T cells at 21 days post-tumor cell inoculation. (**g**) Frequency of total CD8 T cells in the (left) blood and (right) lungs of WT, KO, and KO mice depleted of CD8 T cells. (**h**) Frequency of total CD4 T cells in the (left) blood and (right) lungs of WT, KO, and KO mice depleted of CD4 T cells. Across all panels, data represent mean ± SEM. All *p*-values computed by ordinary one-way ANOVA *t-*test.

